## Supplementary Materials for "An afferent white matter pathway from the pulvinar to the amygdala facilitates fear recognition"

### Dynamic causal modelling

Within the DCM subsample of 237 participants, 49 were unrelated. For this sample, the winning family was the "Dual with SC and PUL input" (expected probability = 64.86%, exceedance probability = 99.92%). The winning model was within this family (expected probability = 21.15%, exceedance probability = 72.77%) and was the same as the winning model from the full 237 participant sample. Classical statistics on the exponentiated parameter estimates showed that all B parameter estimates were significant except the backward connections between right and left FG to IOG, as was found in the full sample.

We conducted the same series of eight correlations between the structural connectivity measures (global track count and summed weights) and effective connectivity (A and B parameter estimates), while also removing 1 outlier. After correcting for multiple comparisons, we found that participants with greater global fibre count along the right PUL-AMG connection also had stronger modulatory connectivity ($r$ = .422, $p_{bonf}$ = .025) along the same connection.

### Full results tables

#### Comparison of tractographically-reconstructed fibres

##### SC-PUL and PUL-AMG

Repeated measures ANOVAs were conducted to compare the fibre counts (global tractography) or apparent fibre density (local tractography) between pathways (SC-PUL, PUL-AMG) and hemispheres (left, right). Absolute relative head motion was included as a covariate of no interest in each test. Greenhouse Geisser corrections made when Mauchly's test of sphericity was significant. All confidence intervals adjusted for Bonferroni correction. Follow-up paired *t*-tests conducted for significant interactions, bootstrapped with 1,000 iterations. Outliers were removed according to whether participants had data on at least one variable with a standardised residual score above or below 3.

***Table 1.*** **Global tractography fibre counts**: 2 (pathway: SC-PUL, PUL-AMG) × 2 (hemisphere: left, right) design. Four outliers removed out of 622.

| [illegible] | **ANOVA Main Effects** | ***t*-tests for Simple Effects** |
|---|---|---|

| | Effect | *df* | *F* | *p* | $\eta_p^2$ | *df* | *t* | *p* |
|---|---|---|---|---|---|---|---|---|
| | * Pathway | (1,616) | 433.286 | $2.842 \times 10^{-73}$ * | 0.413 | | | |
| | * Hemisphere | (1,616) | 7.583 | 0.006 * | 0.012 | | | |
| | * Pathway × Hemisphere | (1,616) | 16.025 | $7.000 \times 10^{-5}$ * | 0.025 | | | |
| | *Left vs. Right SC-PUL* | | | | | 617 | 1.013 | .311 |
| | * *Left vs. Right PUL-AMG* | | | | | 617 | -9.785 | $4.070 \times 10^{-21}$ * |
| | Pathway × Head Motion | (1,616) | 0.313 | .576 | $5.090 \times 10^{-4}$ * | | | |
| | Hemisphere × Head Motion | (1,616) | 0.055 | .813 | $9.100 \times 10^{-5}$ * | | | |
| | Pathway × Hemisphere × Head Motion | (1,616) | 0.332 | .710 | $2.250 \times 10^{-4}$ * | | | |
| **No Outliers Removed** | * Pathway | (1,620) | 32.416 | $2.194 \times 10^{-8}$ * | .049 | | | |
| | * Hemisphere | (1,620) | 9.175 | .003 * | .015 | | | |
| | * Pathway × Hemisphere | (1,620) | 5.807 | .016 * | .009 | | | |
| | *Left vs. Right SC-PUL* | | | | | 621 | 0.654 | .514 |
| | * *Left vs. Right PUL-AMG* | | | | | 621 | -8.904 | $5.906 \times 10^{-18}$ * |
| | Pathway × Head Motion | (1,620) | 0.030 | .863 | $4.800 \times 10^{-5}$ | | | |
| | Hemisphere × Head Motion | (1,620) | 1.155 | .283 | .002 | | | |
| | Pathway × Hemisphere × Head Motion | (1,620) | 0.332 | .565 | .001 | | | |

* $p < .05$

***Table 2.*** **Local tractography fibre counts**: 2 (pathway: SC-PUL, PUL-AMG) × 2 (hemisphere: left, right) design. Thirteen outliers removed out of 622.

| | **ANOVA Main Effects** | | | | | ***t*-tests for Simple Effects** | | |
|---|---|---|---|---|---|---|---|---|
| | Effect | *df* | *F* | *p* | $\eta_p^2$ | *df* | *t* | *p* |
| **Outliers Removed** | * Pathway | (1,607) | 69.586 | $4.930 \times 10^{-16}$ * | 0.103 | | | |
| | Hemisphere | (1,607) | 1.387 | 0.239 | 0.002 | | | |
| | * Pathway × Hemisphere | (1,607) | 162.48 | $3.828 \times 10^{-33}$ * | 0.211 | | | |
| | * *Left vs. Right SC-PUL* | | | | | 608 | 10.749 | $8.596 \times 10^{-25}$ * |
| | * *Left vs. Right PUL-AMG* | | | | | 608 | -18.205 | $1.960 \times 10^{-59}$ * |
| | Pathway × Head Motion | (1,607) | 1.815 | .178 | .003 | | | |
| | Hemisphere × Head Motion | (1,607) | 1.39 | .239 | .002 | | | |
| | Pathway × Hemisphere × Head Motion | (1,607) | 1.425 | .233 | .002 | | | |
| **N** | * Pathway | (1,620) | 59.749 | $4.365 \times 10^{-14}$ * | .088 | | | |

| | | | | | | | |
|---|---|---|---|---|---|---|---|
| Hemisphere | (1,620) | 0.757 | .385 | .001 | | | |
| * Pathway × Hemisphere | (1,620) | 116.962 | $4.316 \times 10^{-25}$ * | .159 | | | |
| ** Left vs. Right SC-PUL* | | | | | 621 | 9.378 | $1.243 \times 10^{-19}$ * |
| ** Left vs. Right PUL-AMG* | | | | | 621 | -17.515 | $4.068 \times 10^{-56}$ * |
| Pathway × Head Motion | (1,620) | 1.222 | .269 | .002 | | | |
| Hemisphere × Head Motion | (1,620) | 1.511 | .219 | .002 | | | |
| Pathway × Hemisphere × Head Motion | (1,620) | 0.617 | .433 | .001 | | | |

* $p < .05$

### Pulvinar and amygdala subregions

Repeated measures ANOVAs were conducted to compare the fibre counts (global tractography) or apparent fibre density (local tractography) on each pathway (SC-PUL, PUL-AMG) independently. Subregions (5 clusters for the pulvinar, 3 subregions for the amygdala) and hemispheres (left, right) were compared. Absolute relative head motion was included as a covariate of no interest in each test. Greenhouse Geisser corrections made when Mauchly’s test of sphericity was significant. All confidence intervals adjusted for Bonferroni correction. Follow-up paired *t*-tests conducted for significant interactions, bootstrapped with 1,000 iterations. Outliers were removed according to whether participants had data on at least one variable with a standardised residual score above or below 3.

***Table 3.*** **Global tractography fibre counts (terminations in pulvinar subregions) for SC-PUL:** 5 (cluster: inferior, medial, anterior, superior, lateral) × 2 (hemisphere: left, right) design. Ten outliers removed out of 622.

| | **ANOVA Main Effects** | | | | | ***t*-tests for Simple Effects** | | |
|---|---|---|---|---|---|---|---|---|
| | **Effect** | ***df*** | ***F*** | ***p*** | $\eta_p^2$ | ***df*** | ***t*** | ***p*** |
| **Outliers Removed** | Hemisphere | (1,610) | 3.817 | .051 | .090 | | | |
| | * Cluster | (3,2060) | 214.015 | $1.584 \times 10^{-133}$ * | .260 | | | |
| | * Hemisphere × Cluster | (3,2014) | 23.624 | $2.870 \times 10^{-16}$ * | .037 | | | |
| | ** Left: Inferior vs. Anterior* | | | | | 611 | 4.820 | $1.815 \times 10^{-6}$ * |
| | ** Left: Anterior vs. Medial* | | | | | 611 | 2.754 | 0.006 * |
| | ** Left: Medial vs. Lateral* | | | | | 611 | 11.635 | $2.114 \times 10^{-28}$ * |
| | ** Left: Lateral vs. Superior* | | | | | 611 | 15.938 | $4.456 \times 10^{-48}$ * |
| | ** Right: Anterior vs. Inferior* | | | | | 611 | 4.228 | $2.715 \times 10^{-5}$ * |
| | ** Right: Inferior vs. Lateral* | | | | | 611 | 13.982 | $9.574 \times 10^{-39}$ * |
| | *Right: Lateral vs. Medial* | | | | | 611 | 2.045 | 0.041 |
| | ** Right: Medial vs. Superior* | | | | | 611 | 21.108 | $1.076 \times 10^{-74}$ * |

| | | | | | | | | |
|---|---|---|---|---|---|---|---|---|
| | Hemisphere × Head Motion | (1,610) | 0.620 | 0.431 | .001 | | | |
| | Cluster × Head Motion | (3,2060) | 0.478 | 0.720 | .001 | | | |
| | Hemisphere × Cluster × Head Motion | (3,2014) | 0.306 | 0.840 | .001 | | | |
| **No Outliers Removed** | * Hemisphere | (1,620) | 5.749 | .017 * | .009 | | | |
| | * Cluster | (1,913) | 52.174 | $2.998 \times 10^{-17}$ * | .078 | | | |
| | * Hemisphere × Cluster | (3,1743) | 16.591 | $4.129 \times 10^{-10}$ * | .026 | | | |
| | ** Left: Inferior vs. Anterior* | | | | | 621 | 4.191 | $3.200 \times 10^{-5}$ * |
| | *Left: Anterior vs. Medial* | | | | | 621 | 2.314 | .021 |
| | ** Left: Medial vs. Lateral* | | | | | 621 | 6.385 | $3.361 \times 10^{-10}$ * |
| | ** Left: Lateral vs. Superior* | | | | | 621 | 14.235 | $5.357 \times 10^{-40}$ * |
| | ** Right: Anterior vs. Inferior* | | | | | 621 | 3.695 | $2.390 \times 10^{-5}$ * |
| | ** Right: Inferior vs. Lateral* | | | | | 621 | 7.619 | $9.590 \times 10^{-14}$ * |
| | *Right: Lateral vs. Medial* | | | | | 621 | -0.187 | .852 |
| | ** Right: Medial vs. Superior* | | | | | 621 | 12.957 | $3.715 \times 10^{-34}$ * |
| | Hemisphere × Head Motion | (1,620) | 1.555 | .213 | .003 | | | |
| | Cluster × Head Motion | (1,913) | 0.159 | .786 | $2.570 \times 10^{-4}$ | | | |
| | Hemisphere × Cluster × Head Motion | (3,1743) | 0.706 | .539 | .001 | | | |

* $p < .05$ (for *t*-tests, only * if < .05 Bonferroni corrected)

***Table 4.*** **Global tractography fibre counts (terminations in pulvinar subregions) for PUL-AMG:** 5 (cluster: inferior, medial, anterior, superior, lateral) × 2 (hemisphere: left, right) design. Sixty-eight outliers removed out of 622.

| | **ANOVA Main Effects** | | | | | ***t*-tests for Simple Effects** | | |
|---|---|---|---|---|---|---|---|---|
| | **Effect** | ***df*** | ***F*** | ***p*** | $\eta_p^2$ | ***df*** | ***t*** | ***p*** |
| **Outliers Removed** | * Hemisphere | (1,552) | 22.147 | $3.197\times 10^{-6}$ * | .039 | | | |
| | * Cluster | (2,1139) | 354.261 | $2.662 \times 10^{-123}$ * | .391 | | | |
| | * Hemisphere × Cluster | (2,1135) | 15.334 | $1.946 \times 10^{-7}$ * | .027 | | | |
| | ** Left: Inferior vs. Medial* | | | | | 533 | 24.849 | $4.248 \times 10^{-92}$ * |
| | ** Left: Medial vs. Lateral* | | | | | 533 | 4.209 | $2.998 \times 10^{-5}$ * |
| | *Left: Lateral vs. Anterior* | | | | | 533 | 2.458 | .014 |
| | ** Left: Anterior vs. Superior* | | | | | 533 | 4.665 | $3.868 \times 10^{-6}$ * |
| | ** Right: Inferior vs. Lateral* | | | | | 533 | 25.742 | $1.184 \times 10^{-96}$ * |
| | ** Right: Lateral vs. Superior* | | | | | 533 | 7.609 | $1.197 \times 10^{-13}$ * |
| | *Right: Superior vs. Medial* | | | | | 533 | 2.044 | .041 |
| | *Right: Medial vs. Anterior* | | | | | 533 | 4.133 | $4.141 \times 10^{-5}$ * |

| | Effect | df | F | p | $\eta_p^2$ | df | t | p |
|---|---|---|---|---|---|---|---|---|
| | Hemisphere × Head Motion | (1,552) | 1.067 | .302 | .002 | | | |
| | Cluster × Head Motion | (2,1139) | 1.885 | .151 | .003 | | | |
| | Hemisphere × Cluster × Head Motion | (2,1135) | 0.601 | .553 | .001 | | | |
| **No Outliers Removed** | * Hemisphere | (1,620) | 33.464 | $1.152 \times 10^{-8}$ * | .051 | | | |
| | * Cluster | (2,1153) | 320.406 | $3.145 \times 10^{-105}$ * | .341 | | | |
| | * Hemisphere × Cluster | (2,1390) | 14.935 | $9.810 \times 10^{-8}$ * | .024 | | | |
| | ** Left: Inferior vs. Anterior* | | | | | 621 | 23.865 | $8.002 \times 10^{-90}$ * |
| | *Left: Anterior vs. Medial* | | | | | 621 | 3.099 | .002 |
| | ** Left: Medial vs. Lateral* | | | | | 621 | 3.021 | .003 |
| | ** Left: Lateral vs. Superior* | | | | | 621 | 3.835 | $1.380 \times 10^{-4}$ * |
| | ** Right: Anterior vs. Inferior* | | | | | 621 | 25.424 | $2.827 \times 10^{-98}$ * |
| | ** Right: Inferior vs. Lateral* | | | | | 621 | 7.909 | $1.190 \times 10^{-14}$ * |
| | *Right: Lateral vs. Medial* | | | | | 621 | 0.657 | .511 |
| | ** Right: Medial vs. Superior* | | | | | 621 | 3.925 | $9.600 \times 10^{-5}$ * |
| | Hemisphere × Head Motion | (1,620) | 0.278 | .598 | $4.490 \times 10^{-4}$ | | | |
| | Cluster × Head Motion | (2,1153) | 1.134 | .319 | .002 | | | |
| | Hemisphere × Cluster × Head Motion | (2,1390) | 0.512 | .620 | .001 | | | |

* $p < .05$ (for *t*-tests, only * if < .05 Bonferroni corrected)

***Table 5.*** **Global tractography fibre counts (terminations in amygdala subregions) for PUL-AMG:** 3 (cluster: centromedial, basolateral, superficial) × 2 (hemisphere: left, right) design. Sixteen outliers removed.

| | **ANOVA Main Effects** | | | | | ***t*-tests for Simple Effects** | | |
|---|---|---|---|---|---|---|---|---|
| | **Effect** | ***df*** | ***F*** | ***p*** | $\eta_p^2$ | ***df*** | ***t*** | ***p*** |
| **Outliers Removed** | Hemisphere | (1,604) | 1.419 | .234 | .002 | | | |
| | * Cluster | (2,1161) | 80.779 | $1.909 \times 10^{-32}$ * | .118 | | | |
| | * Hemisphere × Cluster | (2,1171) | 28.278 | $2.060 \times 10^{-12}$ * | .045 | | | |
| | ** Left: Basolateral vs. Centromedial* | | | | | 605 | 8.741 | $2.279 \times 10^{-17}$ * |
| | ** Left: Centromedial vs. Superficial* | | | | | 605 | 5.354 | $1.222 \times 10^{-7}$ * |
| | ** Right: Basolateral vs. Superficial* | | | | | 605 | 11.515 | $7.018 \times 10^{-28}$ * |
| | ** Right: Superficial vs. Centromedial* | | | | | 605 | 10.700 | $1.376 \times 10^{-24}$ * |
| | Hemisphere × Head Motion | (1,604) | 0.586 | .444 | $.695 \times 10^{-4}$ | | | |
| | Cluster × Head Motion | (2,1161) | 0.892 | .407 | .001 | | | |
| | Hemisphere × Cluster × Head Motion | (2,1171) | 0.261 | .764 | $4.315 \times 10^{-4}$ | | | |

| | Effect | df | F | p | $\eta_p^2$ | t | p |
|---|---|---|---|---|---|---|---|
| **No Outliers Removed** | Hemisphere | (1,620) | 2.295 | .130 | .004 | | |
| | * Cluster | (2,1212) | 79.253 | 2.017 × 10$^{-32}$ * | .113 | | |
| | * Hemisphere × Cluster | (2,1201) | 27.754 | 3.363 × 10$^{-12}$ * | .043 | | |
| | ** Left: Basolateral vs. Centromedial* | | | | 621 | 8.806 | 1.284 × 10$^{-17}$ * |
| | ** Left: Centromedial vs. Superficial* | | | | 621 | 5.119 | 4.098 × 10$^{-7}$ * |
| | ** Right: Basolateral vs. Superficial* | | | | 621 | 11.753 | 6.132 × 10$^{-29}$ * |
| | ** Right: Superficial vs. Centromedial* | | | | 621 | 10.027 | 4.961 × 10$^{-22}$ * |
| | Hemisphere × Head Motion | (1,620) | 0.014 | .907 | 2.200 × 10$^{-4}$ | | |
| | Cluster × Head Motion | (2,1212) | 0.515 | .593 | .001 | | |
| | Hemisphere × Cluster × Head Motion | (2,1201) | 0.164 | .842 | 2.650 × 10$^{-4}$ | | |

* $p < .05$ (for *t*-tests, only * if < .05 Bonferroni corrected)

***Table 6.*** **Local tractography fibre counts (terminations in pulvinar subregions) for SC-PUL:** 5 (cluster: inferior, medial, anterior, superior, lateral) × 2 (hemisphere: left, right) design. Sixty-seven outliers removed out of 622.

| | **ANOVA Main Effects** | | | | | ***t*-tests for Simple Effects** | | |
|---|---|---|---|---|---|---|---|---|
| | **Effect** | ***df*** | ***F*** | ***p*** | $\eta_p^2$ | ***df*** | ***t*** | ***p*** |
| **Outliers Removed** | * Hemisphere | (1,553) | 23.577 | 1.564 × 10$^{-6}$ * | .041 | | | |
| | * Cluster | (2,1010) | 873.922 | 6.337 × 10$^{-209}$ * | .612 | | | |
| | * Hemisphere × Cluster | (2,1176) | 64.370 | 1.002 × 10$^{-28}$ * | .104 | | | |
| | ** Left: Anterior vs. Inferior* | | | | | 554 | 22.373 | 1.709 × 10$^{-79}$ * |
| | ** Left: Inferior vs. Medial* | | | | | 554 | 9.742 | 8.353 × 10$^{-21}$ * |
| | ** Left: Medial vs. Lateral* | | | | | 554 | 30.710 | 1.089 × 10$^{-121}$ * |
| | ** Left: Lateral vs. Superior* | | | | | 554 | 16.514 | 4.103 × 10$^{-50}$ * |
| | ** Right: Anterior vs. Inferior* | | | | | 554 | 39.241 | 4.436 × 10$^{-162}$ * |
| | ** Right: Inferior vs. Medial* | | | | | 554 | 9.157 | 1.022 × 10$^{-18}$ * |
| | ** Right: Medial vs. Lateral* | | | | | 554 | 25.287 | 2.203 × 10$^{-94}$ * |
| | ** Right: Lateral vs. Superior* | | | | | 554 | 15.213 | 6.282 × 10$^{-44}$ * |
| | Hemisphere × Head Motion | (1,553) | 0.358 | 0.550 | .001 | | | |
| | Cluster × Head Motion | (2,1010) | 0.495 | 0.593 | .001 | | | |
| | Hemisphere × Cluster × Head Motion | (2,1176) | 0.604 | 0.557 | .001 | | | |
| **No Outliers Removed** | * Hemisphere | (1,620) | 23.600 | 2.000 × 10$^{-6}$ * | .037 | | | |
| | * Cluster | (2,1244) | 831.764 | 1.817 × 10$^{-230}$ * | .573 | | | |
| | * Hemisphere × Cluster | (2,1354) | 69.262 | 1.107 × 10$^{-31}$ * | .100 | | | |
| | ** Left: Anterior vs. Inferior* | | | | | 621 | 22.745 | 9.069 × 10$^{-84}$ * |
| | ** Left: Inferior vs. Medial* | | | | | 621 | 8.436 | 2.312 × 10$^{-16}$ * |

| | Effect | df | F | p | $\eta_p^2$ | df | t | p |
|---|---|---|---|---|---|---|---|---|
| | ** Left: Medial vs. Lateral* | | | | | 621 | 29.562 | $1.448 \times 10^{-120}$ * |
| | ** Left: Lateral vs. Superior* | | | | | 621 | 13.832 | $3.972 \times 10^{-39}$ * |
| | ** Right: Anterior vs. Inferior* | | | | | 621 | 39.100 | $1.326 \times 10^{-169}$ * |
| | ** Right: Inferior vs. Medial* | | | | | 621 | 7.436 | $3.463 \times 10^{-13}$ * |
| | ** Right: Medial vs. Lateral* | | | | | 621 | 23.544 | $4.380 \times 10^{-88}$ * |
| | ** Right: Lateral vs. Superior* | | | | | 621 | 12.964 | $3.461 \times 10^{-34}$ * |
| | Hemisphere × Head Motion | (1,620) | 0.161 | .688 | $2.60 \times 10^{-4}$ | | | |
| | Cluster × Head Motion | (2,1244) | 0.830 | .437 | .001 | | | |
| | Hemisphere × Cluster × Head Motion | (2,1354) | 1.023 | .365 | .002 | | | |

* $p < .05$ (for $t$-tests, only * if < .05 Bonferroni corrected)

***Table 7.*** **Local tractography fibre counts (terminations in pulvinar subregions) for AMG-PUL:** 5 (cluster: inferior, medial, anterior, superior, lateral) × 2 (hemisphere: left, right) design. Seventy-eight outliers removed out of 622.

| | ANOVA Main Effects | | | | | *t*-tests for Simple Effects | | |
|---|---|---|---|---|---|---|---|---|
| | **Effect** | ***df*** | ***F*** | ***p*** | $\eta_p^2$ | ***df*** | ***t*** | ***p*** |
| **Outliers Removed** | * Hemisphere | (1,542) | 108.301 | $3.019 \times 10^{-23}$ * | .167 | | | |
| | * Cluster | (1,544) | 2159.827 | $9.415 \times 10^{-192}$ * | .799 | | | |
| | * Hemisphere × Cluster | (1,544) | 105.772 | $7.093 \times 10^{-23}$ * | .163 | | | |
| | ** Left: Inferior vs. Lateral* | | | | | 543 | 57.694 | $8.996 \times 10^{-234}$ * |
| | ** Left: Lateral vs. Medial* | | | | | 543 | 10.633 | $4.080 \times 10^{-24}$ * |
| | ** Left: Medial vs. Anterior* | | | | | 543 | 12.744 | $9.913 \times 10^{-33}$ * |
| | ** Left: Anterior vs. Superior* | | | | | 543 | 5.897 | $6.503 \times 10^{-9}$ * |
| | ** Right: Inferior vs. Lateral* | | | | | 543 | 59.079 | $1.332 \times 10^{-238}$ * |
| | ** Right: Lateral vs. Superior* | | | | | 543 | 27.277 | $7.886 \times 10^{-104}$ * |
| | ** Right: Superior vs. Anterior* | | | | | 543 | 14.006 | $2.839 \times 10^{-38}$ * |
| | *Right: Anterior vs. Medial* | | | | | 543 | 0.815 | .416 |
| | Hemisphere × Head Motion | (1,542) | 4.749 | .030 * | 0.009 | | | |
| | Cluster × Head Motion | (1,544) | 19.569 | $1.146 \times 10^{-5}$ * | 0.035 | | | |
| | Hemisphere × Cluster × Head Motion | (1,544) | 4.689 | .031 * | 0.009 | | | |
| **No Outliers Removed** | * Hemisphere | (1,620) | 110.587 | $6.523 \times 10^{-24}$ * | .151 | | | |
| | * Cluster | (1,622) | 2139.649 | $4.887 \times 10^{-204}$ * | .775 | | | |
| | * Hemisphere × Cluster | (1,624) | 107.010 | $2.219 \times 10^{-23}$ * | .147 | | | |
| | ** Left: Inferior vs. Lateral* | | | | | 621 | 57.120 | $2.174 \times 10^{-249}$ * |
| | ** Left: Lateral vs. Medial* | | | | | 621 | 10.169 | $1.427 \times 10^{-22}$ * |

| | | | | | | | |
|---|---|---|---|---|---|---|---|
| ** Left: Medial vs. Anterior* | | | | | 621 | 10.609 | $2.812 \times 10^{-24}$ * |
| ** Left: Anterior vs. Superior* | | | | | 621 | 5.153 | $3.446 \times 10^{-7}$ * |
| ** Right: Inferior vs. Lateral* | | | | | 621 | 61.529 | $1.963 \times 10^{-266}$ * |
| ** Right: Lateral vs. Superior* | | | | | 621 | 24.241 | $7.327 \times 10^{-92}$ * |
| *Right: Superior vs. Anterior* | | | | | 621 | 10.891 | $2.122 \times 10^{-25}$ * |
| *Right: Anterior vs. Medial* | | | | | 621 | 0.227 | .821 |
| Hemisphere × Head Motion | (1,620) | 4.775 | .029 * | .147 | | | |
| Cluster × Head Motion | (1,622) | 19.739 | $1.000 \times 10^{-5}$ * | .031 | | | |
| Hemisphere × Cluster × Head Motion | (1,624) | 4.568 | .033 | .007 | | | |

* $p$ < .05 (for *t*-tests, only * if < .05 Bonferroni corrected)

***Table 8.*** **Local tractography fibre counts (terminations in amygdala subregions) for PUL-AMG:** 3 (cluster: centromedial, basolateral, superficial) × 2 (hemisphere: left, right) design. Thirty-seven outliers removed out of 622.

| | **ANOVA Main Effects** | | | | | ***t*-tests for Simple Effects** | | |
|---|---|---|---|---|---|---|---|---|
| | **Effect** | ***df*** | ***F*** | ***p*** | $\eta_p^2$ | ***df*** | ***t*** | ***p*** |
| **Outliers Removed** | Hemisphere | (1,620) | 511.278 | $5.154 \times 10^{-83}$ * | .452 | | | |
| | * Cluster | (2,989) | 308.079 | $7.593 \times 10^{-88}$ * | .332 | | | |
| | * Hemisphere × Cluster | (2,1201) | 27.754 | $3.363 \times 10^{-12}$ * | .043 | | | |
| | ** Left: Basolateral vs. Centremedial* | | | | | 621 | 10.374 | $2.323 \times 10^{-23}$ * |
| | ** Left: Centromedial vs. Superficial* | | | | | 621 | 19.573 | $8.248 \times 10^{-67}$ * |
| | ** Right: Centromedial vs. Basolateral* | | | | | 621 | 25.136 | $1.037 \times 10^{-96}$ * |
| | ** Right: Basolateral vs. Superficial* | | | | | 621 | 27.037 | $5.308 \times 10^{-107}$ * |
| | Hemisphere × Head Motion | (1,620) | 296.624 | .015 * | .009 | | | |
| | Cluster × Head Motion | (2,989) | 0.050 | $4.774 \times 10^{-84}$ | .324 | | | |
| | Hemisphere × Cluster × Head Motion | (2,1201) | 0.917 | .380 | .001 | | | |
| **No Outliers Removed** | Hemisphere | (1,620) | 511.278 | $5.154 \times 10^{-83}$ * | .452 | | | |
| | * Cluster | (2,989) | 308.079 | $7.593 \times 10^{-88}$ * | .332 | | | |
| | * Hemisphere × Cluster | (2,1201) | 27.754 | $3.363 \times 10^{-12}$ * | .043 | | | |
| | ** Left: Basolateral vs. Centremedial* | | | | | 621 | 10.374 | $2.323 \times 10^{-23}$ * |
| | ** Left: Centromedial vs. Superficial* | | | | | 621 | 19.573 | $8.248 \times 10^{-67}$ * |
| | ** Right: Centromedial vs. Basolateral* | | | | | 621 | 25.136 | $1.037 \times 10^{-96}$ * |
| | ** Right: Basolateral vs. Superficial* | | | | | 621 | 27.037 | $5.308 \times 10^{-107}$ * |
| | Hemisphere × Head Motion | (1,620) | 296.624 | .015 * | .009 | | | |
| | Cluster × Head Motion | (2,989) | 0.050 | $4.774 \times 10^{-84}$ | .324 | | | |

| | | | | |
|---|---|---|---|---|
| Hemisphere × Cluster × Head Motion | (2,1201) | 0.917 | .380 | .001 |

* $p < .05$ (for *t*-tests, only * if < .05 Bonferroni corrected)

#### Null tractography comparison

We conducted a series of paired *t*-tests between the number of streamlines generated by local probabilistic tractography using the iFOD2 algorithm vs. the null distribution algorithm implemented in MRtrix 3. Outliers with a difference more than 3 standard deviations from the mean were removed from the 622-participant sample for each paired *t*-test. Bootstrapping was set to 1,000 iterations.

***Table 9*. Paired *t*-tests between local streamline count and null distribution.**

| Comparison | *N* | *M* | *SEM* | *95% CI* | *t* | *df* | *p* |
|---|---|---|---|---|---|---|---|
| Left SC-PUL | 618 | 1384.876 | 16.704 | [1352.073, 1417.68] | 82.907 | 617 | $< 1 \times 10^{-243}$ * |
| Right SC-PUL | 618 | -499.555 | 16.430 | [-531.821, -467.289] | -30.404 | 617 | $8.832 \times 10^{-125}$ * |
| Left PUL-AMG | 618 | 351.669 | 8.535 | [334.908, 368.431] | 41.202 | 617 | $2.722 \times 10^{-179}$ * |
| Right PUL-AMG | 620 | 577.535 | 10.385 | [557.142, 597.929] | 55.614 | 619 | $6.091 \times 10^{-243}$ * |

* $p < .05$

### Multivariate diffusion-behaviour relationships

We conducted two separate multivariate analyses of covariance, one for global and one for local tractography measures of fibres, to examine the relationship between emotional expression recognition (fearful, sad, and angry expressions) and fibre density along the subcortical route.

***Table 10.* Global tractography multivariate GLM**. Fibre counts for left and right SC-PUL and PUL-AMG entered as dependent variables. Recognition scores (out of eight) for fearful, sad, and angry expressions entered as covariates of interest. Head motion entered as covariate of no interest. Bootstrapping at 1,000 iterations. Four outliers removed with residuals more than 3 standard deviations from the mean.

| | **Multivariate Tests** | | | | | |
|---|---|---|---|---|---|---|
| | **Effect** | **Wilk's Λ** | ***df*** | ***F*** | ***p*** | $\eta_p^2$ |
| **Outliers Removed** | Fearful | .995 | (4, 609) | 0.715 | .582 | .005 |
| | Sad | .994 | (4, 609) | 0.915 | .455 | .006 |
| | Angry | .998 | (4, 609) | 0.294 | .882 | .002 |
| | Head Motion | .999 | (4, 609) | 0.223 | .926 | .001 |

| Parameter Estimates | | | | | | |
|---|---|---|---|---|---|---|
| **Dependent Variable** | **Parameter** | ***B*** | ***t*** | ***p*** | $\eta_p^2$ | ***95% CI*** |
| *Left SC-PUL* | Fearful | -0.350 | -1.583 | .114 | .004 | [-0.784, 0.084] |
| | Sad | -0.312 | -1.520 | .129 | .004 | [-0.715, 0.091] |
| | Angry | -0.157 | -0.687 | .493 | .001 | [-0.607, 0.292] |
| | Head Motion | -0.359 | -0.677 | .498 | .001 | [-1.401, 0.682] |
| *Right SC-PUL* | Fearful | -0.208 | -0.933 | .351 | .001 | [-0.645, 0.23] |
| | Sad | -0.042 | -0.202 | .840 | $6.700 \times 10^{-5}$ | [-0.448, 0.364] |
| | Angry | -0.046 | -0.197 | .844 | $6.400 \times 10^{-5}$ | [-0.499, 0.408] |
| | Head Motion | -0.268 | -0.501 | .616 | $4.110 \times 10^{-4}$ | [-1.317, 0.781] |
| *Left PUL-AMG* | Fearful | -0.006 | -0.051 | .959 | $4.000 \times 10^{-6}$ | [-0.225, 0.213] |
| | Sad | -0.010 | -0.099 | .921 | $1.600 \times 10^{-5}$ | [-0.213, 0.193] |
| | Angry | -0.024 | -0.205 | .837 | $6.900 \times 10^{-5}$ | [-0.251, 0.203] |
| | Head Motion | 0.103 | 0.386 | .700 | $2.430 \times 10^{-4}$ | [-0.422, 0.628] |
| *Left PUL-AMG* | Fearful | -0.071 | -0.611 | .541 | .001 | [-0.297, 0.156] |
| | Sad | 0.101 | 0.941 | .347 | .001 | [-0.11, 0.311] |
| | Angry | -0.101 | -0.846 | .398 | .001 | [-0.336, 0.134] |
| | Head Motion | -0.103 | -0.373 | .709 | $2.280 \times 10^{-4}$ | [-0.647, 0.44] |

**No Outliers Removed**

| Multivariate Tests | | | | | |
|---|---|---|---|---|---|
| **Effect** | **Wilk's Λ** | ***df*** | ***F*** | ***p*** | $\eta_p^2$ |
| Fearful | .998 | (4,613) | 0.349 | .845 | .002 |
| Sad | .994 | (4,613) | 0.957 | .430 | .006 |
| Angry | .996 | (4,613) | 0.569 | .685 | .004 |
| Head Motion | .998 | (4,613) | 0.287 | .886 | .002 |

| Parameter Estimates | | | | | | |
|---|---|---|---|---|---|---|
| **Dependent Variable** | **Parameter** | ***B*** | ***t*** | ***p*** | $\eta_p^2$ | ***95% CI*** |
| *Left SC-PUL* | Fearful | -0.794 | -0.809 | .419 | .001 | [-2.723, 1.134] |
| | Sad | 0.053 | 0.058 | .954 | $5.000 \times 10^{-7}$ | [-1.736, 1.842] |
| | Angry | 0.057 | 0.056 | .955 | $5.000 \times 10^{-7}$ | [-1.938, 2.052] |

| | | | | | | | |
|---|---|---|---|---|---|---|---|
| | | Head Motion | -0.147 | -0.062 | .950 | 6.000 × $10^{-7}$ | [-4.763, 4.47] |
| | *Right SC-PUL* | Fearful | -0.612 | -0.605 | .546 | .001 | [-2.597, 1.374] |
| | | Sad | 0.482 | 0.514 | .607 | 4.290 × $10^{-4}$ | [-1.36, 2.325] |
| | | Angry | 0.314 | 0.3 | .764 | 1.460 × $10^{-4}$ | [-1.74, 2.369] |
| | | Head Motion | -0.703 | -0.29 | .772 | 1.370 × $10^{-4}$ | [-5.458, 4.052] |
| | *Left PUL-AMG* | Fearful | -0.072 | -0.433 | .665 | 3.050 × $10^{-4}$ | [-0.4, 0.255] |
| | | Sad | 0.018 | 0.117 | .907 | 2.200 × $10^{-5}$ | [-0.286, 0.322] |
| | | Angry | -0.109 | -0.631 | .529 | .001 | [-0.448, 0.23] |
| | | Head Motion | 0.191 | 0.479 | .632 | 3.720 × $10^{-4}$ | [-0.593, 0.976] |
| | *Left PUL-AMG* | Fearful | -0.126 | -0.871 | .384 | .001 | [-0.41, 0.158] |
| | | Sad | 0.12 | 0.894 | .372 | .001 | [-0.144, 0.384] |
| | | Angry | -0.135 | -0.905 | .366 | .001 | [-0.429, 0.158] |
| | | Head Motion | -0.012 | -0.033 | .973 | 2.000 × $10^{-6}$ | [-0.692, 0.669] |

***Table 11.* Local tractography multivariate GLM**. Average apparent fibre density for left and right SC-PUL and PUL-AMG entered as dependent variables. Recognition scores (out of eight) for fearful, sad, and angry expressions entered as covariates of interest. Head motion entered as covariate of no interest. Bootstrapping at 1,000 iterations. Fifteen outliers removed with residuals more than 3 standard deviations from the mean.

| | **Multivariate Tests** | | | | | | |
|---|---|---|---|---|---|---|---|
| | **Effect** | | **Wilk's Λ** | ***df*** | ***F*** | ***p*** | **$\eta_p^2$** |
| **Outliers Removed** | * Fearful | | .984 | (4,598) | 2.501 | .042 * | .016 |
| | Sad | | .991 | (4,598) | 1.363 | .245 | .009 |
| | Angry | | .993 | (4,598) | 1.111 | .350 | .007 |
| | * Head Motion | | .977 | (4,598) | 3.451 | .008 * | .023 |
| | **Parameter Estimates** | | | | | | |
| | **Dependent Variable** | **Parameter** | ***B*** | ***t*** | ***p*** | **$\eta_p^2$** | ***95% CI*** |
| | *Left SC-PUL* | Fearful | 0 | -0.003 | 0.998 | 1.093 × $10^{-8}$ | [-0.163, 0.162] |

| | | | | | | | |
|---|---|---|---|---|---|---|---|
| | | Sad | 0.084 | 1.099 | 0.272 | 0.002 | [-0.066, 0.235] |
| | | Angry | -0.078 | -0.901 | 0.368 | 0.001 | [-0.249, 0.093] |
| | | Head Motion | -0.135 | -0.681 | 0.496 | 0.001 | [-0.526, 0.255] |
| | *Right SC-PUL* | Fearful | -0.046 | -0.645 | 0.519 | 0.001 | [-0.185, 0.094] |
| | | Sad | -0.008 | -0.123 | 0.902 | $2.500 \times 10^{-5}$ | [-0.137, 0.121] |
| | | Angry | 0.043 | 0.574 | 0.566 | 0.001 | [-0.104, 0.189] |
| | | Head Motion | -0.146 | -0.859 | 0.39 | 0.001 | [-0.48, 0.188] |
| | *Left PUL-AMG* | * Fearful | 0.14 | 2.887 | 0.004 * | 0.014 | [0.045, 0.235] |
| | | Sad | -0.084 | -1.855 | 0.064 | 0.006 | [-0.172, 0.005] |
| | | Angry | -0.052 | -1.018 | 0.309 | 0.002 | [-0.152, 0.048] |
| | | Head Motion | -0.21 | -1.802 | 0.072 | 0.005 | [-0.438, 0.019] |
| | *Right PUL-AMG* | * Fearful | 0.143 | 2.523 | 0.012 * | 0.010 | [0.032, 0.255] |
| | | Sad | -0.042 | -0.787 | 0.431 | 0.001 | [-0.145, 0.062] |
| | | Angry | 0.041 | 0.686 | 0.493 | 0.001 | [-0.076, 0.158] |
| | | * Head Motion | -0.484 | -3.55 | $4.150 \times 10^{-4}$ * | 0.021 | [-0.752, -0.216] |
| **No Outliers Removed** | **Multivariate Tests** | | | | | | |
| | **Effect** | | **Wilk's Λ** | ***df*** | ***F*** | ***p*** | $\eta_p^2$ |
| | * Fearful | | .983 | (4,613) | 2.669 | .031 * | .017 |
| | Sad | | .993 | (4,613) | 1.080 | .366 | .007 |
| | Angry | | .997 | (4,613) | 0.499 | .736 | .003 |
| | * Head Motion | | .982 | (4,613) | 2.792 | .026 * | .018 |
| | **Parameter Estimates** | | | | | | |
| | **Dependent Variable** | **Parameter** | **B** | **t** | **p** | $\eta_p^2$ | **95% CI** |
| | *Left SC-PUL* | Fearful | -0.006 | -0.051 | .960 | $4.000 \times 10^{-6}$ | [-0.231, 0.22] |
| | | Sad | 0.121 | 1.138 | .255 | .002 | [-0.088, 0.33] |
| | | Angry | -0.080 | -0.674 | .500 | .001 | [-0.313, 0.153] |
| | | Head Motion | -0.063 | -0.229 | .819 | $8.500 \times 10^{-5}$ | [-0.602, 0.477] |
| | *Right SC-PUL* | Fearful | -0.097 | -1.037 | .300 | .002 | [-0.28, 0.086] |
| | | Sad | 0.017 | 0.197 | .844 | $6.300 \times 10^{-5}$ | [-0.153, 0.187] |

| | | | | | | | |
|---|---|---|---|---|---|---|---|
| | | Angry | 0.023 | 0.233 | .816 | 8.800 × $10^{-5}$ | [-0.167, 0.212] |
| | | Head Motion | -0.165 | -0.738 | .461 | .001 | [-0.604, 0.274] |
| | *Left PUL-AMG* | * Fearful | 0.137 | 2.650 | .008 * | .011 | [0.036, 0.239] |
| | | Sad | -0.075 | -1.563 | .119 | .004 | [-0.17, 0.019] |
| | | Angry | -0.049 | -0.904 | .366 | .001 | [-0.154, 0.057] |
| | | Head Motion | -0.193 | -1.554 | .121 | .004 | [-0.437, 0.051] |
| | *Left PUL-AMG* | * Fearful | 0.137 | 2.334 | .020 * | .009 | [0.022, 0.253] |
| | | Sad | -0.053 | -0.976 | .330 | .002 | [-0.16, 0.054] |
| | | Angry | 0.001 | 0.019 | .985 | 6.126 × $10^{-7}$ | [-0.118, 0.121] |
| | | * Head Motion | -0.465 | -3.303 | .001 * | .017 | [-0.741, -0.189] |

### Partial correlations between fibre density and effective connectivity parameters

We conducted a series of right-sided Pearson's partial correlations between the fibre density of each connection (left and right SC-PUL and PUL-AMG connections, global and local tractography – giving eight in total) and its corresponding DCM parameter estimate (modulatory effect of Faces > Shapes over region coupling). Head motion was entered as a control variable. Multivariate outliers were detected according to Mahalanobis distance ($df$ = 8, $\chi^2$ criterion = 15.507, $p$ = .05), resulting in 24 participants being excluded from the analysis (N = 213). Note that outliers were substantially influencing the results (see table below for comparison).

**Table 12. Partial correlations between fibre density of each pathway and the corresponding DCM parameter estimate.** Scatterplots show the fibre density (*x*-axis) residuals (after regressing against head motion) and the DCM parameter estimate (*y*-axis) residuals (after regressing against head motion). * $p$ < .05, Bonferroni-corrected

| o | **Global Tractography** |
|---|---|

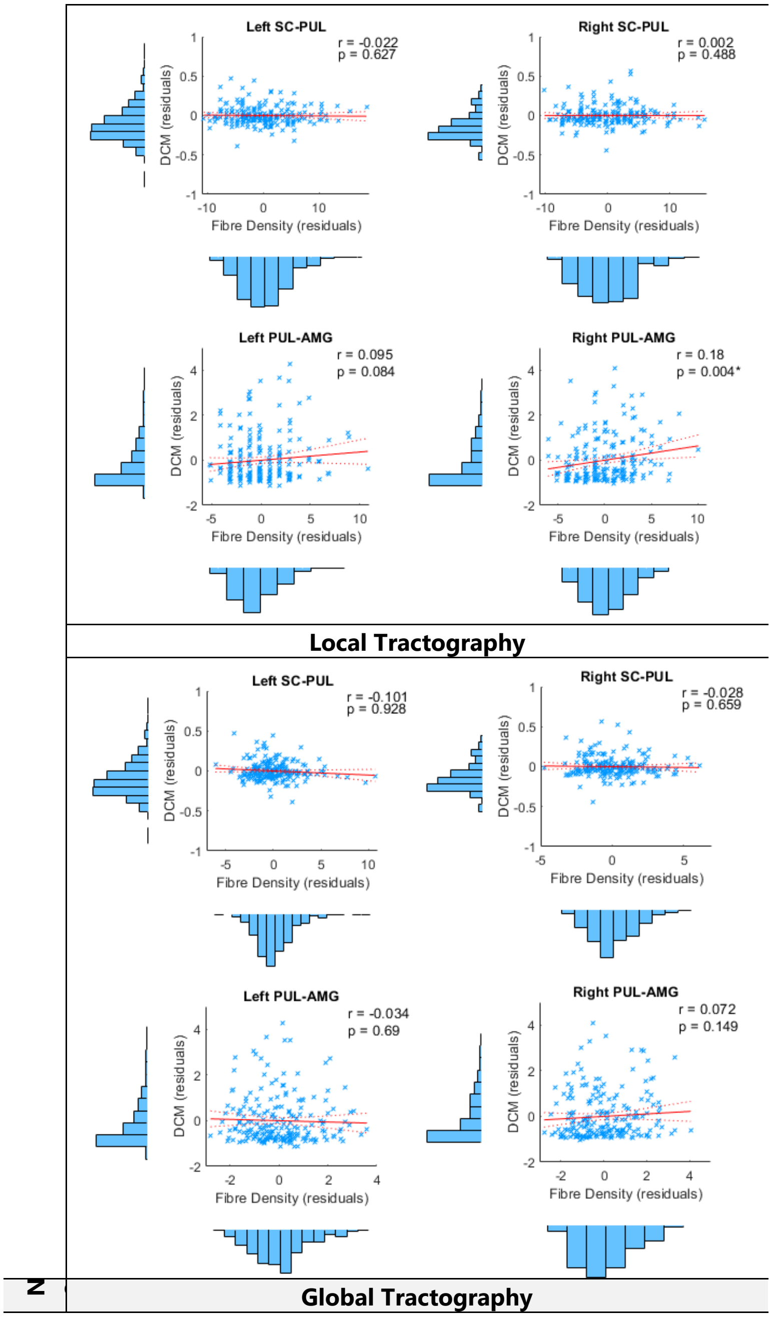
Left SC-PUL
r = -0.022
p = 0.627
Right SC-PUL
r = 0.002
p = 0.488
Left PUL-AMG
r = 0.095
p = 0.084
Right PUL-AMG
r = 0.18
p = 0.004*
DCM (residuals)
Fibre Density (residuals)
Local Tractography
Left SC-PUL
r = -0.101
p = 0.928
Right SC-PUL
r = -0.028
p = 0.659
Left PUL-AMG
r = -0.034
p = 0.69
Right PUL-AMG
r = 0.072
p = 0.149
DCM (residuals)
Fibre Density (residuals)
Global Tractography

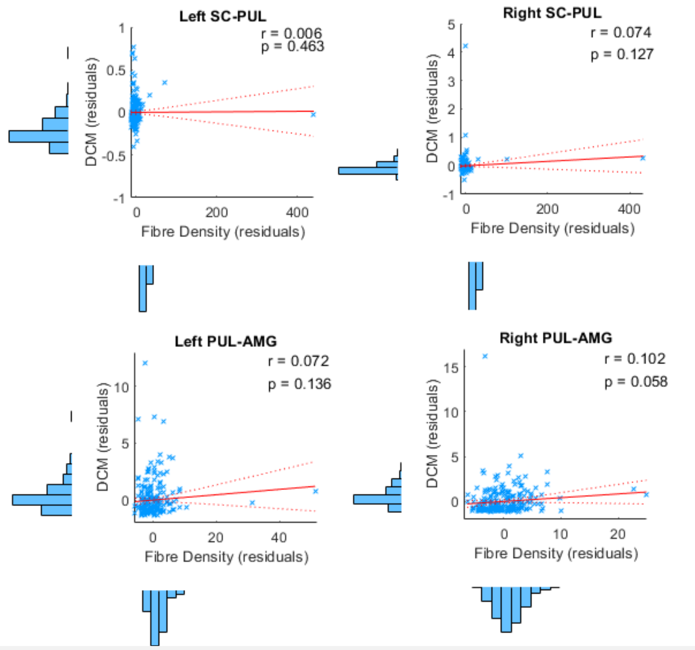

**Local Tractography**

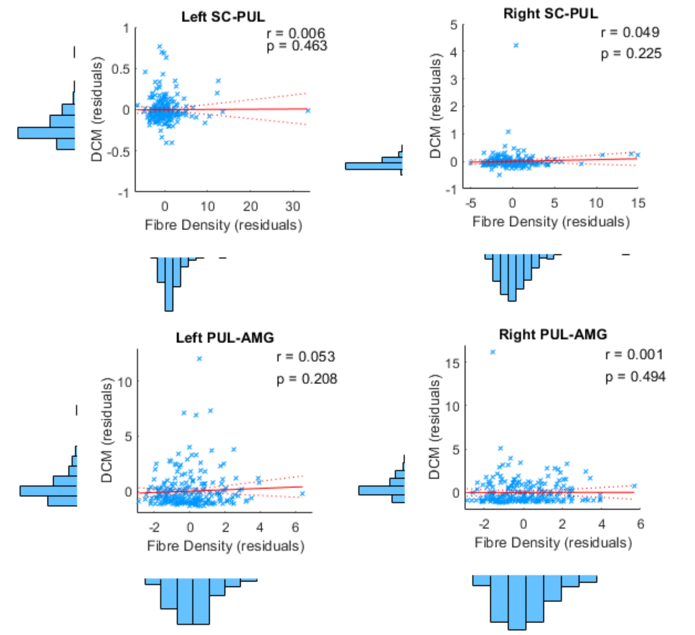

108

### DCM parameter estimates

We conducted one-sample *t*-tests on the exponentiated DCM parameter estimates (B matrix) against a test value of 1 to examine whether modulatory connection strength was consistently greater than the prior within our sample of participants. Outliers were removed that were more than 3 SDs from the mean of each variable (note that this did not change the pattern of results).

**Table 13. Significance of DCM parameter estimates**.

| | Parameter | *M* | *SD* | *df* | *t* | *p* | 95% CI |
|---|---|---|---|---|---|---|---|
| **Outliers Removed** | * Left SC-PUL | 1.063 | 0.115 | 229 | 8.285 | < .001 | [1.048, 1.078] |
| | * Right SC-PUL | 1.059 | 0.124 | 233 | 7.295 | < .001 | [1.043, 1.075] |
| | * Left PUL-AMG | 1.834 | 1.000 | 233 | 12.753 | < .001 | [1.705, 1.962] |
| | * Right PUL-AMG | 1.731 | 0.765 | 230 | 14.514 | < .001 | [1.632, 1.83] |
| | * Left IOG-FG | 1.685 | 0.715 | 228 | 14.485 | < .001 | [1.592, 1.778] |
| | * Right IOG-FG | 2.415 | 2.140 | 233 | 10.111 | < .001 | [2.139, 2.69] |
| | * Left FG-AMG | 1.213 | 0.258 | 233 | 12.614 | < .001 | [1.18, 1.246] |
| | * Right FG-AMG | 1.190 | 0.217 | 230 | 13.308 | < .001 | [1.162, 1.218] |
| | Left FG-IOG | 0.988 | 0.211 | 230 | -0.842 | 0.401 | [0.961, 1.016] |
| | Right FG-IOG | 1.037 | 0.270 | 231 | 2.095 | 0.037 | [1.002, 1.072] |
| | * Left AMG-FG | 1.813 | 1.330 | 231 | 9.306 | < .001 | [1.641, 1.985] |
| | * Right AMG-FG | 1.878 | 1.101 | 228 | 12.078 | < .001 | [1.735, 2.022] |
| **No Outliers Removed** | **Parameter** | ***M*** | ***SD*** | ***df*** | ***t*** | ***p*** | **95% CI** |
| | * Left SC-PUL | 1.066 | 0.157 | 236 | 6.481 | < .001 | [1.046, 1.086] |
| | * Right SC-PUL | 1.068 | 0.190 | 236 | 5.510 | < .001 | [1.044, 1.093] |
| | * Left PUL-AMG | 1.916 | 1.235 | 236 | 11.415 | < .001 | [1.758, 2.074] |
| | * Right PUL-AMG | 1.821 | 0.944 | 236 | 13.382 | < .001 | [1.7, 1.942] |
| | * Left IOG-FG | 1.864 | 1.214 | 236 | 10.957 | < .001 | [1.709, 2.02] |
| | * Right IOG-FG | 2.772 | 4.249 | 236 | 6.422 | < .001 | [2.229, 3.316] |
| | * Left FG-AMG | 1.251 | 0.446 | 236 | 8.661 | < .001 | [1.194, 1.308] |
| | * Right FG-AMG | 1.218 | 0.278 | 236 | 12.081 | < .001 | [1.182, 1.253] |
| | Left FG-IOG | 1.028 | 0.329 | 236 | 1.311 | 0.191 | [0.986, 1.07] |
| | Right FG-IOG | 1.069 | 0.348 | 236 | 3.071 | 0.002 | [1.025, 1.114] |
| | * Left AMG-FG | 2.057 | 2.166 | 236 | 7.517 | < .001 | [1.78, 2.335] |
| | * Right AMG-FG | 2.092 | 1.596 | 236 | 10.536 | < .001 | [1.888, 2.296] |

* $p < .001$

### VOI inclusion/exclusion criteria

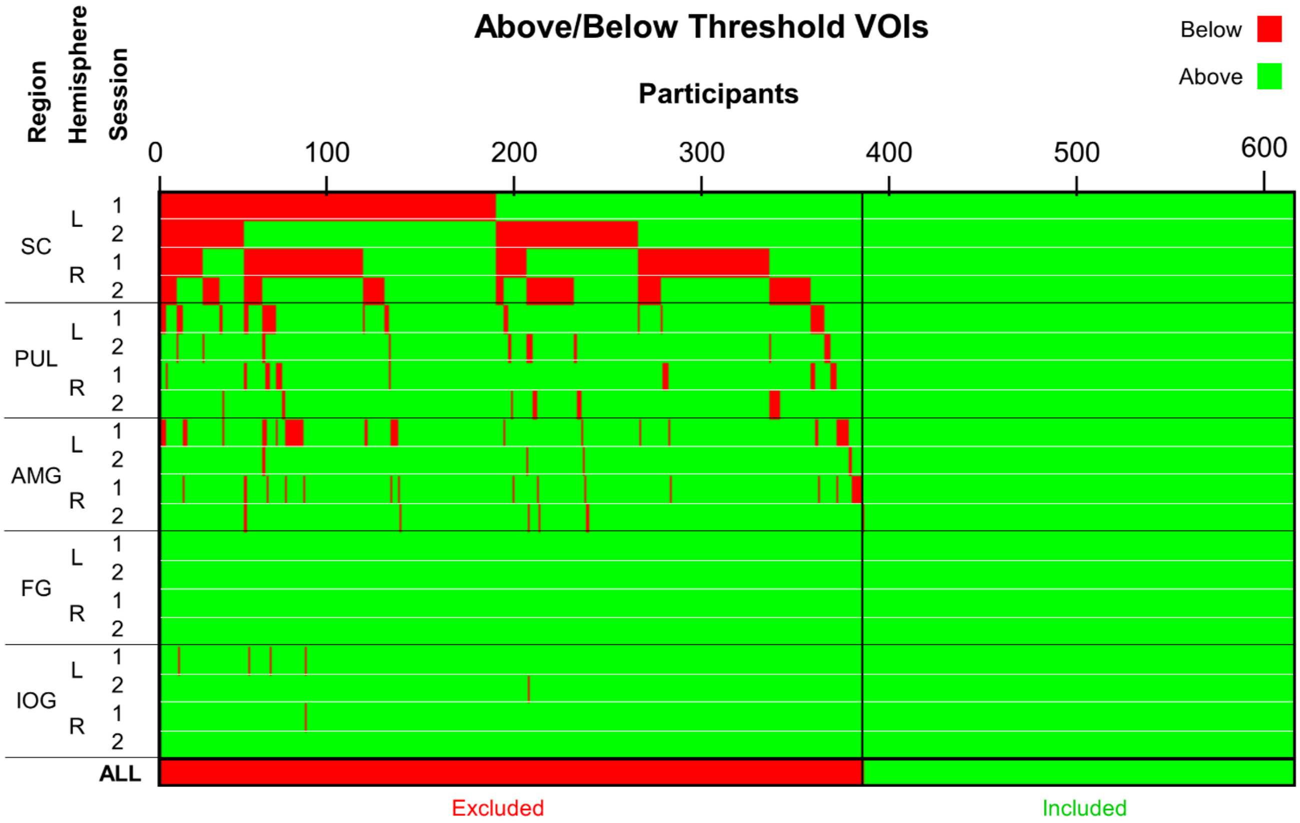

**Figure 4 – Figure Supplement 1. Participants with above or below threshold fMRI signal in all VOIs for inclusion into DCM stage.** Threshold was set at $p < .05$ uncorrected, using the [Faces – Shapes] contrast. Each column is a participant (N = 622) and each row is a different region (SC = superior colliculus, PUL = pulvinar, AMG = amygdala, FG = fusiform gyrus, IOG = inferior occipital gyrus) with 2 sessions each (i.e. each fMRI run) per hemisphere (L = left, R = right). Red indicates below-threshold signal and green indicates above-threshold signal. The bottom row ('ALL') indicates whether the participants were included or excluded from further DCM analysis, based on whether they had any below-threshold VOIs.
